# A Common Class of Transcripts with 5′-Intron Depletion, Distinct Early Coding Sequence Features, and N^1^-Methyladenosine Modification

**DOI:** 10.1101/057455

**Authors:** Can Cenik, Hon Nian Chua, Guramrit Singh, Abdalla Akef, Michael P Snyder, Alexander F. Palazzo, Melissa J Moore, Frederick P Roth

## Abstract

Introns are found in 5’ untranslated regions (5’UTRs) for 35% of all human transcripts. These 5’UTR introns are not randomly distributed: genes that encode secreted, membrane-bound and mitochondrial proteins are less likely to have them. Curiously, transcripts lacking 5’UTR introns tend to harbor specific RNA sequence elements in their early coding regions. To model and understand the connection between coding-region sequence and 5’UTR intron status, we developed a classifier that can predict 5’UTR intron status with >80% accuracy using only sequence features in the early coding region. Thus, the classifier identifies transcripts with <u>5</u>’ proximal-<u>i</u>ntron-<u>m</u>inus-like-coding regions (“5IM” transcripts). Unexpectedly, we found that the early coding sequence features defining 5IM transcripts are widespread, appearing in 21% of all human RefSeq transcripts. The 5IM class of transcripts is enriched for non-AUG start codons, more extensive secondary structure both preceding the start codon and near the 5’ cap, greater dependence on eIF4E for translation, and association with ER-proximal ribosomes. 5IM transcripts are bound by the Exon Junction Complex (EJC) at non-canonical 5’ proximal positions. Finally, N^1^-methyladenosines are specifically enriched in the early coding regions of 5IM transcripts. Taken together, our analyses point to the existence of a distinct 5IM class comprising ∼20% of human transcripts. This class is defined by depletion of 5’ proximal introns, presence of specific RNA sequence features associated with low translation efficiency, N^1^-methyladenosines in the early coding region, and enrichment for non-canonical binding by the Exon Junction Complex.

## Introduction

Approximately 35% of all human transcripts harbor introns in their 5’ untranslated regions (5’UTRs) (Cenik et al. 2010; Hong et al. 2006). Among genes with 5’UTR introns (5UIs), those annotated as “regulatory” are significantly overrepresented, while there is an under-representation of genes encoding proteins that are targeted to either the endoplasmic reticulum (ER) or mitochondria (Cenik et al. 2011). For transcripts that encode ER- and mitochondria-targeted proteins, 5UI depletion is associated with presence of specific RNA sequences (Cenik et al. 2011; Palazzo et al. 2007, 2013). Specifically, nuclear export of an otherwise inefficiently exported microinjected mRNA or cDNA transcript can be promoted by an ER-targeting signal sequence-containing region (SSCRs) or mitochondrial signal sequence coding region (MSCRs) from a gene lacking 5’ UTR introns (Cenik et al. 2011; Lee et al. 2015). However, more recent studies suggest that many SSCRs have little impact on nuclear export for RNAs transcribed *in vivo* (Lee et al. 2015), but rather enhance translation in a RanBP2-dependent manner (Mahadevan et al. 2013).

Among SSCR- and MSCR-containing transcripts (referred to hereafter as SSCR and MSCR transcripts), ∼75% lack 5’ UTR introns (“5UI^−^” transcripts) and ∼25% have them (“5UI^+^” transcripts). These two groups have markedly different sequence compositions at the 5’ ends of their coding sequences. 5UI^−^ transcripts tend to have lower adenine content (Palazzo et al. 2007) and use codons with fewer uracils and adenines than 5UI^+^ transcripts (Cenik et al. 2011). Their signal sequences also contain leucine and arginine more often than the biochemically similar amino acids isoleucine and lysine, respectively. Leucine and arginine codons contain fewer adenine and thymine nucleotides, consistent with adenine and thymine depletion. This depletion is also associated with the presence of a specific GC-rich RNA motif in the early coding region of 5UI^−^ transcripts (Cenik et al. 2011).

Despite some knowledge as to their early coding region features, key questions about this class of 5UI^−^ transcripts have remained unanswered: Do the above sequence features extend beyond SSCR- and MSCR-containing transcripts to other 5UI^−^ genes? Do 5UI^−^ transcripts having these features share common functional or regulatory features? What binding factor(s) recognize these RNA elements? A more complete model of the relationship of early coding features and 5UI^−^ status would begin to address these questions.

Here, to better understand the relationship between early coding region features and 5UI status, we undertook an integrative machine learning approach. We reasoned that a machine learning classifier which could identify 5UI^−^ transcripts solely from early coding sequence would potentially provide two types of insight: First, it could systematically identify predictive features. Second, the subset of 5UI^−^ transcripts which could be identified by the classifier might then represent a functionally distinct transcript class. Having developed such a classifier, we found that it identified ∼21% of all human transcripts as harboring coding regions characteristic of 5UI^−^ transcripts. While many of these transcripts encode ER- and mitochondrial-targeted proteins, many others encode nuclear and cytoplasmic proteins. This class of transcripts shares characteristics such as a tendency to lack 5’ proximal introns, to contain non-canonical Exon Junction Complex binding sites, to have multiple features associated with lower intrinsic translation efficiency, and to have an increased incidence of N^1-^ methyladenosine modification.

## Results

### A classifier that predicts 5UI status using only early coding sequence information

To better understand the previously-reported enigmatic relationship between certain early coding region sequences and the absence of a 5UI, we sought to model this relationship. Specifically, we used a random forest classifier (Breiman 2001) to learn the relationship between 5UI absence and a collection of 36 different nucleotide-level features extracted from the first 99 nucleotides of all human coding regions (CDS) (**Figs 1A-C; Table S1**; Methods). We then used all transcripts known to contain an SSCR (a total of 3743 transcripts clusters; Methods), regardless of 5UI status, as our training set. This training constraint ensured that all input nucleotide sequences were subject to similar functional constraints at the protein level. Thus, we sought to identify sequence features that differ between 5UI^−^ and 5UI^+^ transcripts at the RNA level.

**Fig 1.**
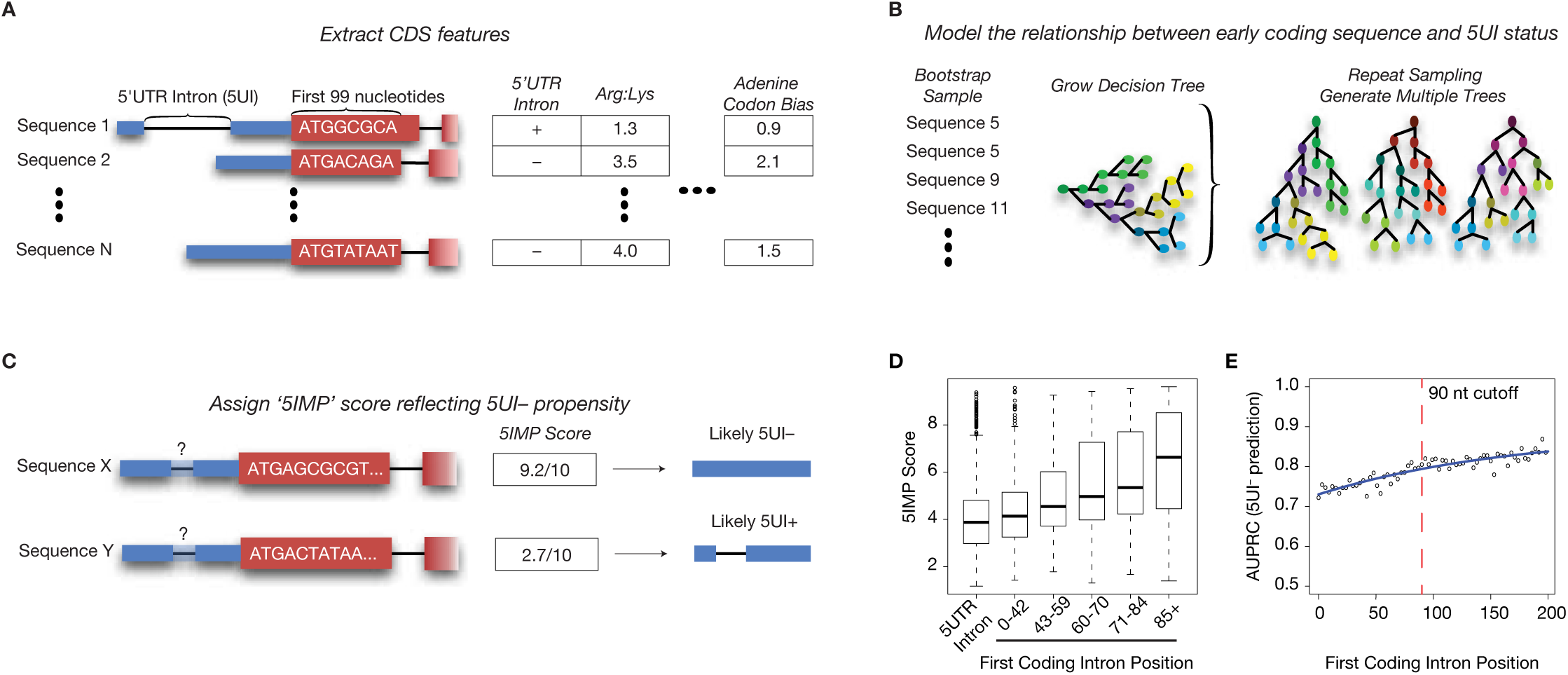
Modeling the relationship between sequence features in the early coding region and the absence of 5’UTR introns (5UIs). **(a)** For all human transcripts, information about 36 nucleotide-level features of the early coding region (first 99 nucleotides) and 5UI presence was extracted. **(b)** Transcripts containing a signal sequence coding region (SSCR) were used to train a Random Forest classifier that modeled the relationship between 5UI absence and 36 sequence features. **(c)** With this classifier, all human transcripts were assigned a score that quantifies the likelihood of 5UI absence based on specific RNA sequence features in the early coding region. Transcripts with high scores are thus considered to have 5’-proximal intron minus-like coding regions (5IMs). “5’UTR-intron-minus-predictor” (5IMP) score distributions for SSCR-containing transcripts shift to higher scores with later-appearing first introns, suggesting that 5IM coding region features not only predict lack of a 5UI, but also lack of early coding region introns. **(e)** Classifier performance was optimized by excluding 5UI^−^ transcripts with introns appearing early in the coding region. Cross-validation performance (area under the precision recall curve, AUPRC) was examined for a series of alternative 5IM classifiers using different first-intron-position criterion for excluding 5UI^−^ transcripts from the training set (Methods).

Our classifier assigns to each transcript a “5’UTR-intron-minus-predictor” (5IMP) score between 0 and 10, where higher scores correspond to a higher likelihood of being 5UI^−^ (**Fig 1C**). Interestingly, preliminary ranking of the 5UI^−^ transcripts by 5IMP score revealed a relationship between the position of the first intron in the coding region and the 5IMP score. 5UI^−^ transcripts for which the first intron was more than 85 nts downstream of the start codon had the highest 5IMP scores. Furthermore, the closer the first intron was to the start codon, the lower the 5IMP score (**Fig 1D**). We explored this relationship further by training classifiers that increasingly excluded from the training set 5UI^−^ transcripts according to the distance of the first intron from the 5’ end of the coding region. This revealed that classifier performance, as measured by the area under the precision recall curve (AUPRC), increased as a function of the distance from start codon to first intron distance (Methods, **Fig 1E**). Thus, the RNA sequence features we identified as being predictive of 5UI^−^ transcripts are more accurately described as being predictors of transcripts without 5’-proximal introns.

To minimize the impact of transcripts that may ‘behave’ as though they were 5UI^+^ due to an intron early in the coding region, we eliminated 5UI^−^ SSCR transcripts with a first intron <90 nts downstream of the start codon (Methods) and generated a new classifier. Discriminative motif features were learned independently (Methods), and performance of this new classifier was gauged using 10-fold cross validation. We assessed cross-validation performance in two ways: 1) in terms of the area under the receiver operating curve (AUC)— which can be thought of as a measure of average recall across a range of false positive rates; 2) in terms of area under the precision vs recall curve (AUPRC), which can be thought of as the average precision (fraction of predictions which are correct) across a range of recall values. Specifically, the classifier showed an AUC of 74% and AUPRC of 88% (**Fig 2A, yellow curves**; exceeding AUC 50% and AUPRC 71%, the performance value expected of a naïve predictor). We used this optimized classifier for all subsequent analyses.

**Fig 2.**
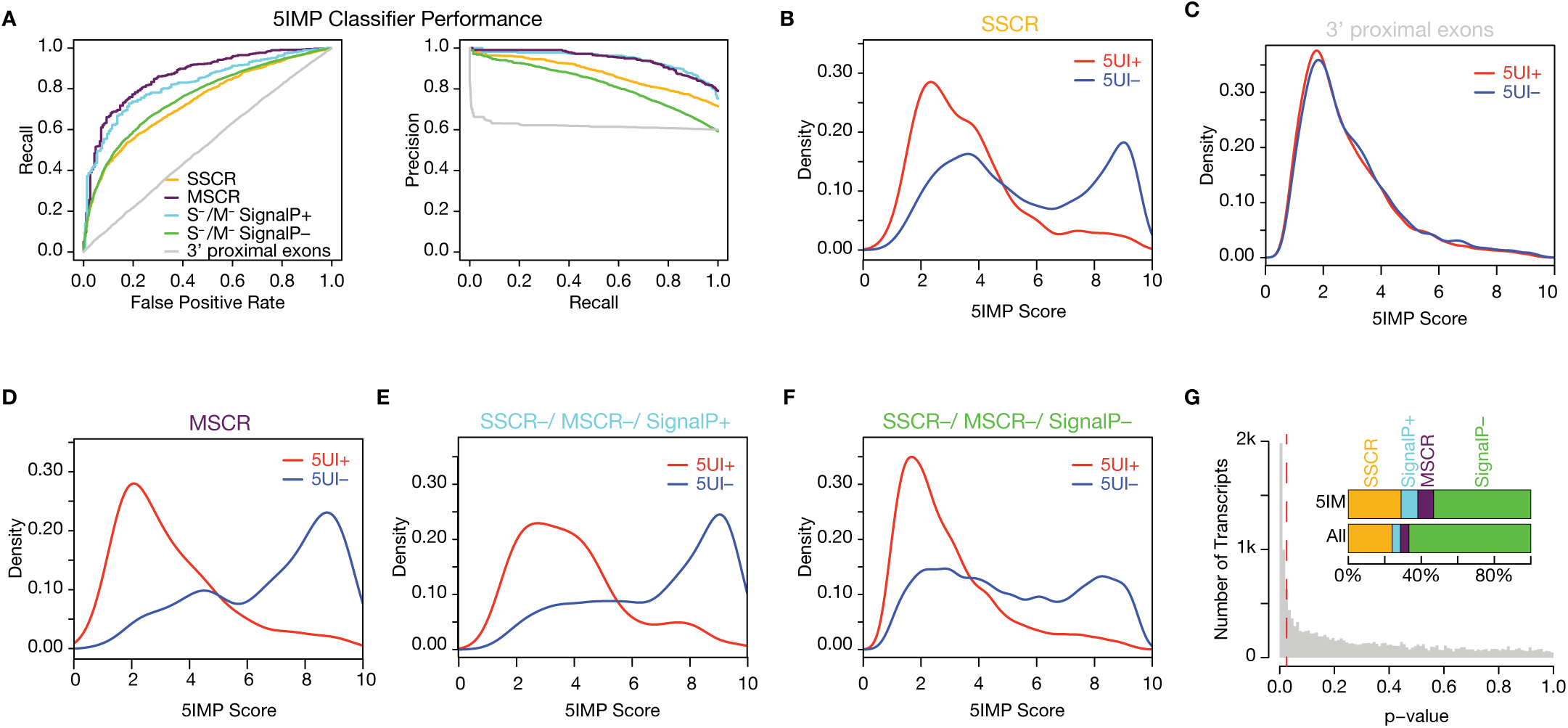
Predicting 5UI status accurately using only early coding sequences. **(a)** As judged by area under the receiver operating characteristic curve (AUROC) and AUPRC, The 5IM classifier performed well for several different transcript classes. **(b)** The distribution of 5IMP scores reveals clear separation of 5UI^+^ and 5UI^−^ transcripts for SSCR-containing transcripts, where each SSCR-containing transcript was scored using a classifier that did not use that transcript in training (Methods). **(c)** Coding sequence features that are predictive of 5’ proximal intron presence are restricted to the early coding region. This was supported by identical 5IM classifier score distributions with respect to 5UI presence for negative control sequences, each derived from a single randomly chosen ‘window’ downstream of the 3rd exon from one of the evaluated transcripts. **(d)** MSCR transcripts exhibited a major difference in 5IMP scores based on their 5UI status even though no MSCR transcripts were used in training the classifier. **(e)** Transcripts predicted to contain signal peptides (SignalP^+^) had a 5IMP score distribution similar to that of SSCR-containing transcripts. **(f)** After eliminating SSCR, MSCR, and SignalP^+^ transcripts, the remaining S^−^/MSCR^−^ SignalP^−^ transcripts were still significantly enriched for high 5IM classifier scores among 5UI^−^ transcripts. **(g)** The control set of randomly chosen sequences downstream of the 3^rd^ exon from each transcript was used to calculate an empirical cumulative null distribution of 5IMP scores. Using this function, we determined the p-value corresponding to the 5IMP score for all transcripts. The red dashed line indicates the p-value corresponding to 5% False Discovery Rate. The inset depicts the distribution of various classes of mRNAs among the input set and 5IM transcripts.

SSCR transcripts exhibited markedly different 5IMP score distributions for the 5UI^+^ and 5UI^−^ subsets (**Fig 2B**). The 5UI^+^ score distribution was unimodal with a peak at ∼2.4. In contrast, the 5UI^−^ score distribution was bimodal with one peak at ∼3.6 and another at ∼9, suggesting the existence of at least two underlying 5UI^−^ transcript classes. The peak at score 3.6 resembled the 5UI^+^ peak. Also contributing to the peak at 3.6 is the set of 5UI^−^ transcripts harboring an intron in the first 90 nts of the CDS (55% of all 5UI^−^ transcripts). The other distinct high-scoring 5UI^−^ class (peak at score 9) is composed of transcripts that have specific 5UI^−^-predictive RNA sequence elements within the early coding region.

We next wished to evaluate whether our classifier was discriminating 5UI^+^ and 5UI^−^ SSCR transcripts using signals that appear specifically in the early coding region as opposed to signals that appear broadly across the coding region. To do so, for every transcript we randomly chose 99 nts from the region downstream of the 3^rd^ exon. The 5IMP score distributions of these ‘3’ proximal exon’ sets were identical for 5UI^+^ and 5UI^−^ transcripts (**Figs 2A, and 2C**), confirming that the sequence features that distinguish 5UI^+^ and 5UI^−^ transcripts are specific to the early coding region.

### RNA elements associated with 5UI^−^ transcripts are pervasive in the human genome

Having trained the classifier on SSCR transcripts, we wondered how well it would predict the 5UI status of other transcripts. Despite having been trained exclusively on SSCR transcripts, the classifier performed remarkably well on MSCR transcripts, achieving an AUC of 86% and AUPRC of 95% (**Fig 2A, purple line; Fig 2D**; as compared with 50% and 77%, respectively, expected by chance). This result suggests that RNA elements within early coding regions of 5UI^−^ MSCR-transcripts are similar to those in 5UI^−^ SSCR-transcripts despite distinct functional constraints at the protein level.

We next wondered whether the class of 5UI^−^ transcripts that can be predicted on the basis of early coding region features is restricted to transcripts encoding proteins trafficked to the ER or mitochondria, or is instead a more general class of transcripts. We therefore asked whether the classifier could predict 5UI^−^ status in transcripts that contain neither an SSCR nor an MSCR (“S^−^/M^−^” transcripts). Because unannotated SSCRs could confound this analysis, we first used SignalP 3.0 to identify S^−^/M^−^ transcripts most likely to contain an unannotated SSCR (Bendtsen et al. 2004). These ‘SignalP^+^‘ transcripts had a 5IMP score distribution comparable to those of known SSCR and MSCR transcripts (**Fig 2E**), and the classifier worked well to identify the 5UI^−^ subset of these transcripts (AUC 82% and AUPRC 95%, **Fig 2A, light blue line**). While 5UI^+^ SignalP^+^ transcripts had predominantly low 5IMP scores, 5UI^−^ SignalP^+^ 5IMP scores were strongly skewed towards high 5IMP scores (peak at ∼9; **Fig 2E**). These results were consistent with the idea that SignalP^+^ transcripts do contain many unannotated SSCRs.

Having considered SignalP^+^ transcripts as well as SSCR- and MSCR-containing transcripts, we used the classifier to calculate 5IMP scores for all remaining “S^−^/M^−^/SignalP^−^” transcripts. Although the performance was weaker on this gene set, it was still better than expected of a naive predictor (**Fig 2A, green line**). 5UI^+^ S^−^/M^−^/SignalP^−^ transcripts were strongly skewed toward low 5IMP scores (**Fig 2F**). Surprisingly, however, a significant fraction of 5UI^−^ S^−^/M^−^/SignalP^−^ transcripts had high 5IMP scores (∼18%). Thus, our results suggest a broad class of transcripts with early coding regions carrying sequence signals that predict the absence of a 5’proximal intron, or in other words, a class of transcripts with <u>5</u>’proximal-<u>i</u>ntron-<u>m</u>inus-like coding regions. Hereafter we refer to transcripts in this class as “5IM” transcripts.

We sought to identify what fraction of transcripts have 5IMP scores that exceed what would be expected in the absence of 5UI^−^-predictive early coding region signals. To establish this expectation, we used the above-described negative control set of equal-length coding sequences from outside of the early coding region. By quantifying the excess of high-scoring sequences in the real distribution relative to this control distribution, we estimate that 21% of all human transcripts are 5IM transcripts (a 5IMP score of 7.41 corresponds to a 5% False Discovery Rate; **Fig 2G**). The set of 5IM transcripts defined by our classifier (**Table S2**) includes many that do not encode ER-targeted or mitochondrial proteins. The distribution of various classes of mRNAs among the 5IM transcripts was: 38% ER-targeted (SSCR or SignalP^+^), 9% mitochondrial (MSCR) and 53% other classes (S^−^/M^−^/SignalP^−^) (Figure 2G). These results suggest that RNA-level features prevalent in the early coding regions of 5UI^−^ SSCR and MSCR transcripts are also found in other transcript types (**Figs 2F-G**), and that 5IM transcripts represent a broad class.

### Functional characterization of 5IM transcripts

5IM transcripts are defined by mRNA sequence features. Hence, we hypothesized that 5IM transcripts may be functionally related through shared regulatory mechanisms mediated by the presence of these common features. To this end, we collected large-scale datasets representing diverse attributes covering six broad categories (see Table S3 for a complete list): (1) Curated functional annotations -- e.g., Gene Ontology terms, annotation as a ‘housekeeping’ gene, genes subject to RNA editing; (2) RNA localization -- e.g., to dendrites, to mitochondria; (3) Protein and mRNA half-life, ribosome occupancy and features that decrease stability of one or more mRNA isoforms -- e.g., AU-rich elements (4) Sequence features associated with regulated translation -- e.g., codon optimality, secondary structure near the start codon; (5) Known interactions with RNA-binding proteins or complexes such as Staufen-1, TDP-43, or the Exon Junction Complex (EJC) (6) RNA modifications – i.e., N^1^-methyladenosine (m^1^A).

We adjusted for multiple hypotheses testing at two levels. First, we took a conservative approach (Bonferroni correction) to correct for the number of tested functional characteristics. Second, some of the functional categories were analyzed in more depth and multiple sub-hypotheses were tested within the given category. In this in-depth analysis a false discovery-based correction was adopted. Below, all reported p-values remain significant (padjusted < 0.05) after multiple hypothesis test correction.

No associations between 5IM transcripts and features in categories (1), (2) and (3) were found, other than the already-known enrichments for ER- and mitochondrial-targeted mRNAs. However, analyses for the remaining categories yielded the significant results described below.

### 5IM transcripts have features suggesting lower translation efficiency

Translation regulation is a major determinant of protein levels (Vogel and Marcotte 2012). To investigate potential connections between 5IM transcripts and translational regulation, we examined features associated with translation. Features found to be significant were:

(I) <u>Secondary structures near the start codon</u> can affect initiation rate by modulating start codon recognition (Parsyan et al. 2011). We observed a positive correlation between 5IMP score and the free energy of folding (-ΔG) of the 35 nucleotides immediately preceding the start codon (**Fig 3A**; Spearman rho=0.39; p < 2.2e-16). This suggests that 5IM transcripts have a greater tendency for secondary structure near the start codon, presumably making the start codon less accessible.

**Fig 3.**
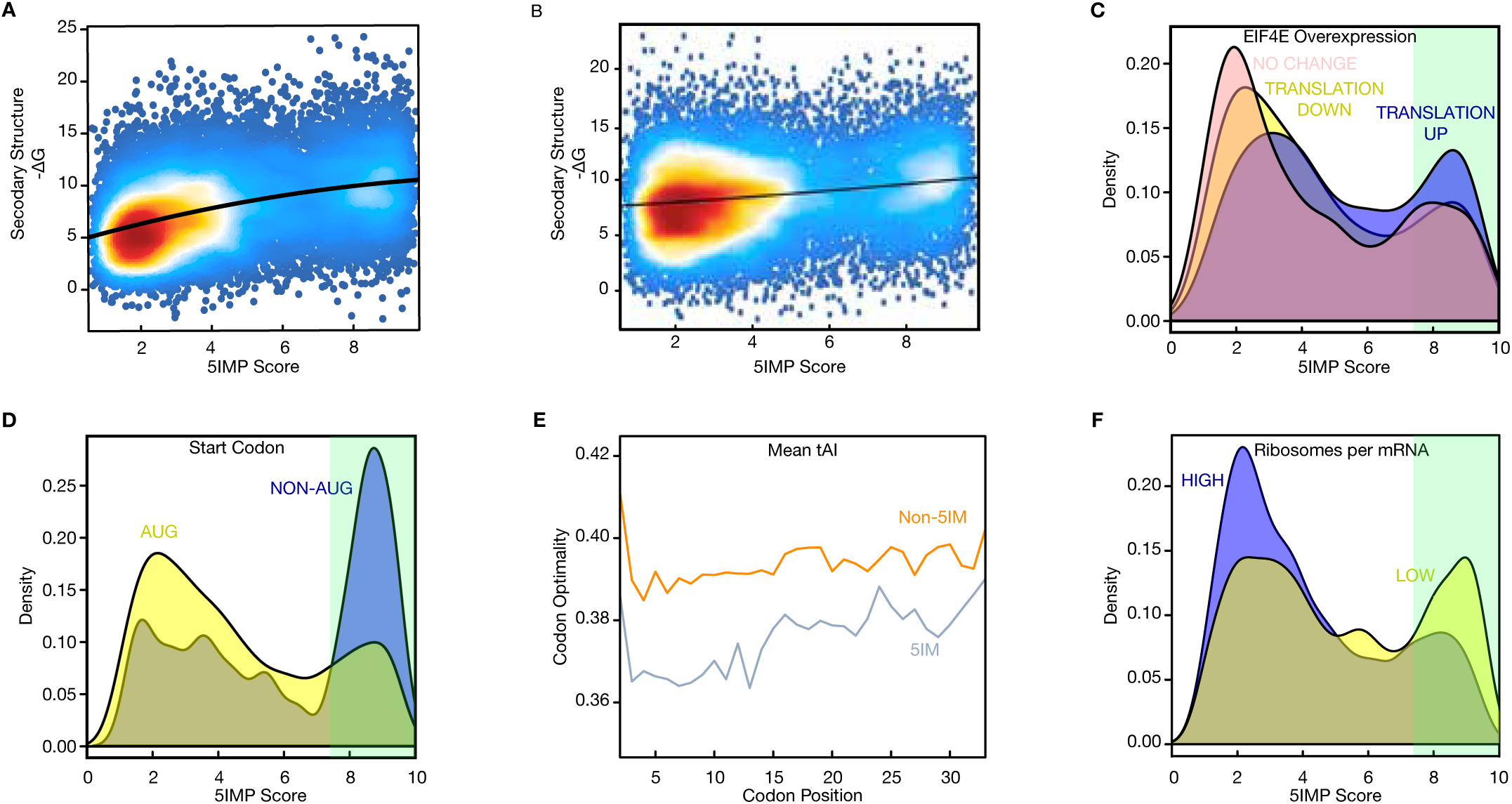
5IM transcripts are more likely to be differentially expressed and have sequence features associated with lower translation efficiency. **(a)** The 5IM classifier score was positively correlated with the propensity for mRNA structure preceding the start codon (−ΔG) (Spearman rho=0.39; p < 2.2e-16). For each transcript, 35 nucleotides immediately upstream of the AUG were used to calculate −ΔG (Methods). **(b)** The 5IM classifier score was positively correlated with the propensity for mRNA structure near the 5’cap (−ΔG) (Spearman rho=0.18; p = 7.9e-130; Methods). **(c)** Transcripts that are translationally upregulated in response to eIF4E overexpression (Larsson et al. 2007) (blue) were enriched for higher 5IMP scores. Light green shading indicates 5IMP scores corresponding to 5% FDR. **(d)** Transcripts with non-AUG start codons (blue) exhibited significantly higher 5IMP scores than transcripts with a canonical ATG start codon (yellow). **(e)** Higher 5IMP scores were associated with less optimal codons (as measured by the tRNA adaptation index, tAI) for the first 33 codons. For all transcripts within each 5IMP score category (blue-high, orange-low), the mean tAI was calculated at each codon position. Start codon was not shown. **(f)** Transcripts with lower translation efficiency were enriched for higher 5IMP scores. Transcripts with translation efficiency one standard deviation below the mean (“LOW” translation, yellow) and one standard deviation higher than the mean (“HIGH” translation, blue) were identified using ribosome profiling and RNA-Seq data from human lymphoblastoid cell lines (Methods).

(II) Similarly, <u>secondary structures near the 5’cap</u> can modulate translation by hindering binding by the 43S-preinitiation complex to the mRNA (Babendure et al. 2006). We observed a positive correlation between 5IMP score and the free energy of folding (-ΔG) of the 5’most 35 nucleotides (**Fig 3B**; Spearman rho=0.18; p = 7.9e-130). This suggests that 5IM transcripts have a greater tendency for secondary structure near the 5’cap, presumably hindering binding by the 43S-preinitiation complex.

(III) <u>eIF4E overexpression</u>. The heterotrimeric translation initiation complex eIF4F (made up of eIF4A, eIF4E and eIF4G) is responsible for facilitating the translation of transcripts with strong 5’UTR secondary structures (Parsyan et al. 2011). The eIF4E subunit binds to the 7mGpppG ‘methyl-G’ cap and the ATP-dependent helicase eIF4A (scaffolded by eIF4G) destabilizes 5’UTR secondary structure (Marintchev et al. 2009). A previous study identified transcripts that were more actively translated under conditions that promote cap-dependent translation (overexpression of eIF4E) (Larsson et al. 2007). In agreement with the observation that 5IM transcripts have more secondary structure upstream of the start codon and near the 5’cap, transcripts with high 5IMP scores were more likely to be translationally, but not transcriptionally, upregulated upon eIF4E overexpression (**Fig 3C**; Wilcoxon Rank Sum Test p = 2.05e-22, and p = 0.28, respectively).

(IV) <u>Non-AUG start codons</u>. Transcripts with non-AUG start codons also have intrinsically low translation initiation efficiencies (Hinnebusch and Lorsch 2012). These mRNAs were greatly enriched among transcripts with high 5IMP scores (Fisher’s Exact Test p = 0.0003; odds ratio = 3.9) and have a median 5IMP score that is 3.57 higher than those with an AUG start (**Fig 3D**).

(V) <u>Codon optimality</u>. The efficiency of translation elongation is affected by codon optimality (Hershberg and Petrov 2008). Although some aspects of this remain controversial (Charneski and Hurst 2013; Shah et al. 2013; Zinshteyn and Gilbert 2013; Gerashchenko and Gladyshev 2015), it is clear that decoding of codons by tRNAs with different abundances can affect the translation rate under conditions of cellular stress (reviewed in Gingold and Pilpel 2011). We therefore examined the tRNA adaptation index (tAI), which correlates with copy numbers of tRNA genes matching a given codon (dos Reis et al. 2004). Specifically, we calculated the median tAI of the first 99 coding nucleotides of each transcript, and found that 5IMP score was negatively correlated with tAI (**Fig S1A**; Spearman Correlation rho= −0.23; p < 2e-16 median tAI and 5IMP score). This effect was restricted to the early coding regions as the negative control set of randomly chosen sequences downstream of the 3^rd^ exon from each transcript did not exhibit a relationship between 5IMP score and codon optimality (**Fig S1B-C**). Thus, 5IM transcripts show reduced codon optimality in early coding regions, suggesting that 5IM transcripts have decreased translation elongation efficiency.

To more precisely determine where the codon optimality phenomenon occurs within the entire early coding region, we grouped transcripts by 5IMP score. For each group, we calculated the mean tAI at codons 2-33 (i.e., nts 4-99). Across this entire region, 5IM transcripts (5IMP >7.41; 5% FDR) had significantly lower tAI values at every codon except codons 24 and 32 (**Fig 3E**; Wilcoxon Rank Sum test Holm-adjusted p < 0.05 for all comparisons). To eliminate potential confounding variables, including nucleotide composition, we performed several additional control analyses (Methods); none of these altered the basic conclusion that 5IM transcripts have lower codon optimality than non-5IM transcripts across the entire early coding region.

(VI) <u>Ribosomes per mRNA</u>. Finally, we examined the relationship between 5IMP score and translation efficiency, as measured by the steady-state number of ribosomes per mRNA molecule. To this end, we used a large dataset of ribosome profiling and RNA-Seq experiments from human lymphoblastoid cell lines (Cenik et al. 2015). From this, we calculated the average number of ribosomes on each transcript and identified transcripts with high or low ribosome occupancy (respectively defined by occupancy at least one standard deviation above or below the mean; see Methods). 5IM transcripts were slightly but significantly depleted in the high ribosome-occupancy category (**Fig 3F**; Fisher’s Exact Test p = 0.0006, odds ratio = 1.3). Moreover, 5IMP scores exhibited a weak but significant negative correlation with the number of ribosomes per mRNA molecule (Spearman rho=-0.11; p = 5.98e-23).

Taken together, all of the above results reveal that 5IM transcripts have sequence features associated with lower translation efficiency, at the stages of both translation initiation and elongation.

### Non-ER trafficked 5IM transcripts are enriched in ER-proximal ribosome occupancy

We next investigated the relationship between 5IMP score and the localization of translation within cells. Exploring the subcellular localization of translation at a transcriptome-scale remains a significant challenge. Yet, a recent study described proximity-specific ribosome profiling to identify mRNAs occupied by ER-proximal ribosomes in both yeast and human cells (Jan et al. 2014). In this method, ribosomes are biotinylated based on their proximity to a marker protein such as Sec61, which localizes to the ER membrane (Jan et al. 2014). For each transcript, the enrichment for biotinylated ribosome occupancy yields a measure of ERproximity of translated mRNAs.

We reanalyzed this dataset to explore the relationship between 5IMP scores and ER-proximal ribosome occupancy in HEK-293 cells. As expected, transcripts that exhibit the highest enrichment for ER-proximal ribosomes were SSCR-containing transcripts and transcripts with other ER-targeting signals. Yet, we noticed a surprising positive correlation between ER-proximal ribosome occupancy and 5IM transcripts with no ER-targeting evidence **(Fig 4**). This relationship was true for both mitochondrial genes (**Fig 4;** Spearman rho=0.43; p < 2.2e-16), and genes with no evidence for either ER- or mitochondrial-targeting (**Fig 4**; Spearman rho=-0.36; p < 2.2e-16). These results suggest that 5IM transcripts are more likely than non-5IM transcripts to engage with ER-proximal ribosomes.

**Fig 4.**
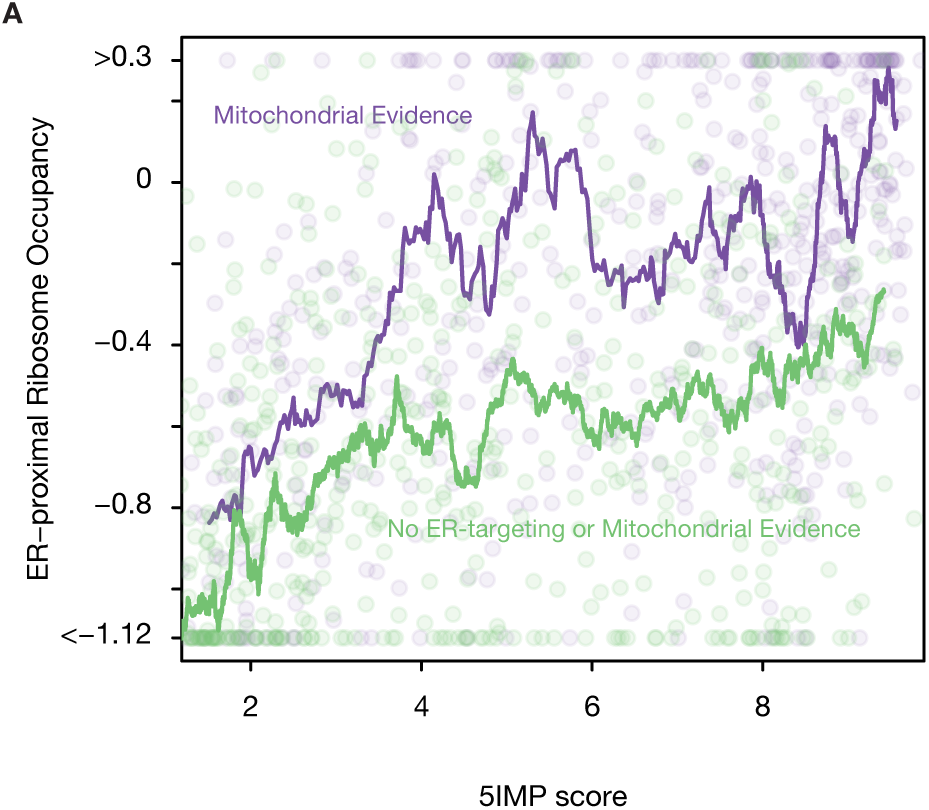
5IM transcripts with no evidence of ER-targeting are more likely to exhibit ER-proximal ribosome occupancy. A moving average of ER-proximal ribosome occupancy was calculated by grouping genes by 5IMP score (see Methods). We plotted the moving average of 5IMP scores for transcripts with no evidence of ER- or mitochondrial targeting (green) or for transcripts predicted to be mitochondrial (purple). We plotted a random subsample of transcripts on top of the moving average (circles).

### 5IM transcripts are strongly enriched in non-canonical EJC occupancy sites

Shared sequence features and functional traits among 5IM transcripts causes one to wonder what common mechanisms might link 5IM sequence features to 5IM traits. For example, 5IM transcripts might share regulation by one or more RNA-binding proteins (RBPs). To investigate this idea further, we tested for enrichment of 5IM transcripts among the experimentally identified targets of 23 different RBPs (including CLIP-Seq, and variants; see Methods). Only one dataset was substantially enriched for high 5IMP scores among targets: a transcriptome-wide map of binding sites of the Exon Junction Complex (EJC) in human cells, obtained via tandem-immunoprecipitation followed by deep sequencing (RIPiT) (Singh et al. 2014, 2012). The EJC is a multi-protein complex that is stably deposited upstream of exon-exon junctions as a consequence of pre-mRNA splicing (Le Hir et al. 2000). RIPiT data confirmed that canonical EJC sites (cEJC sites; sites bound by EJC core factors and appearing ∼24 nts upstream of exon-exon junctions) occupy ∼80% of all possible exon-exon junction sites and are not associated with any sequence motif. Unexpectedly, many EJC-associated footprints outside of the canonical −24 regions were observed (**Fig 5A**) (Singh et al. 2012). These ‘non-canonical’ EJC occupancy sites (ncEJC sites) were associated with multiple sequence motifs, three of which were similar to known recognition motifs for SR proteins that co-purified with the EJC core subunits (Singh et al. 2012). Interestingly, another motif (**Fig 5B**; top) that was specifically found in first exons is not known to be bound by any known RNAbinding protein (Singh et al. 2012). This motif was CG-rich, a sequence feature that also defines 5IM transcripts. This similarity presages the possibility of enrichment of first exon ncEJC sites among 5IM transcripts. Position analysis of called EJC peaks revealed that while only 9% of cEJCs reside in first exons, 19% of all ncEJCs are found there. When we investigated the relationship between 5IMP scores and ncEJCs in early coding regions, we found a striking correspondence—the median 5IMP score was highest for transcripts with the greatest number of ncEJCs (**Fig 5C**; Wilcoxon Rank Sum Test; p < 0.0001). When we repeated this analysis by conditioning on 5UI status, we similarly found that ncEJCs were enriched among transcripts with high 5IMP scores regardless of 5UI status (Fisher’s Exact Test, p < 3.16e-14, odds ratio > 2.3; **Fig 5D**). These results suggest that the striking enrichment of ncEJC peaks in early coding regions was generally applicable to all transcripts with high 5IMP scores regardless of 5UI presence.

**Fig 5.**
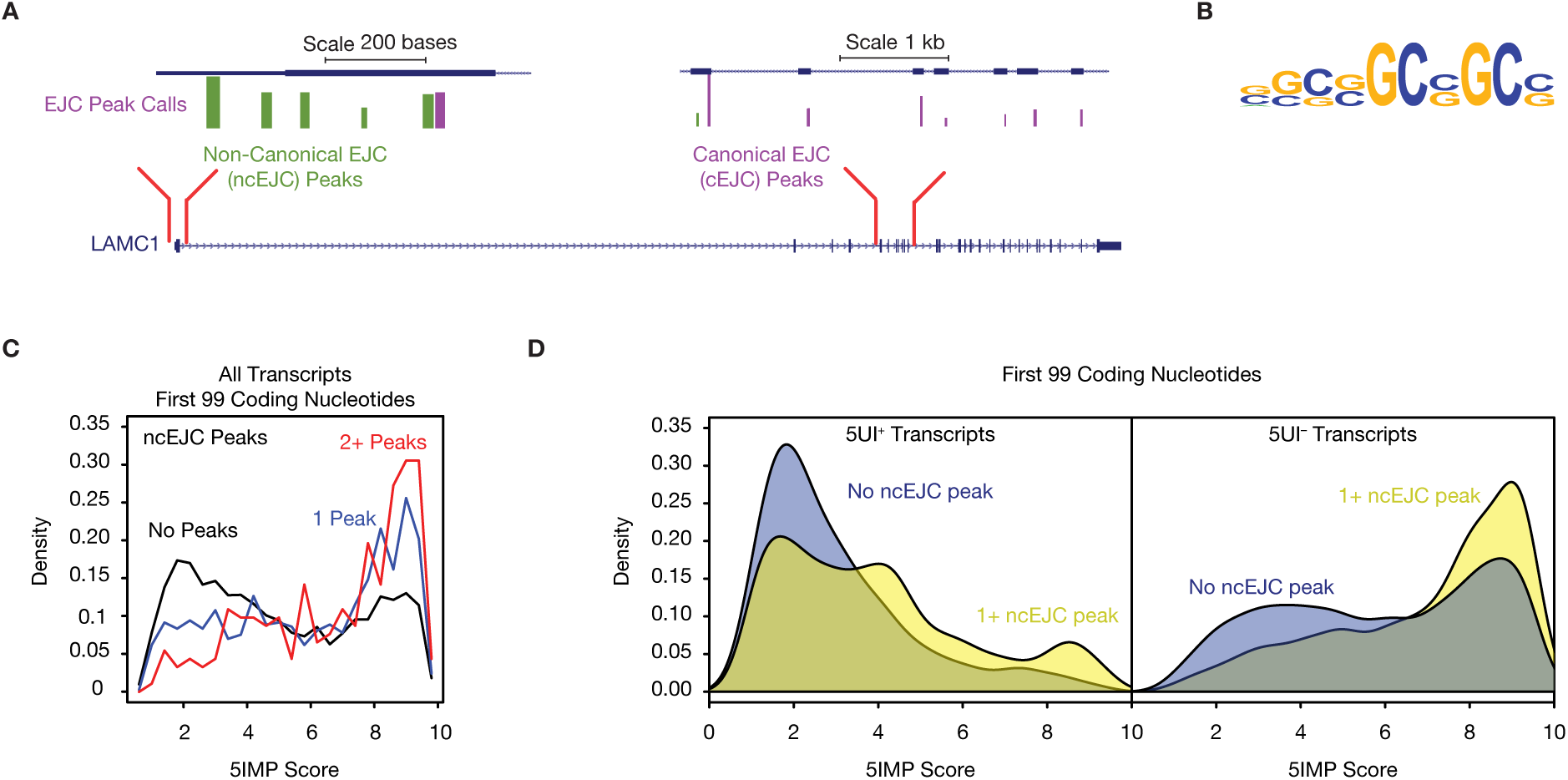
5IM transcripts harbor non-canonical Exon Junction Complex (EJC) binding sites. **(a)** Observed EJC binding sites (Singh et al. 2012) are shown for an example 5IM transcript (*LAMC1*). Canonical EJC binding sites (purple) are ∼24nt upstream of an exon-intron boundary. The remaining binding sites are considered to be non-canonical (green). **(b)** A CG-rich sequence motif previously identified to be enriched among ncEJC binding sites in first exons (Singh et al. 2012) is shown **(c)** 5IMP score for transcripts with zero, one, two or more non-canonical EJC binding sites in the first 99 coding nucleotides reveals that transcripts with high 5IMP scores frequently harbor non-canonical EJC binding sites. **(d)** Transcripts with high 5IMP scores are enriched for non-canonical EJCs regardless of 5UI presence or absence.

### Transcripts harboring N^1^-methyladenosine (m1A) have high 5IMP scores

It is increasingly clear that ribonucleotide base modifications in mRNAs are highly prevalent and can be a mechanism for post-transcriptional regulation (Frye et al. 2016). One RNA modification present towards the 5’ ends of mRNA transcripts is N^1^-methyladenosine (m^1^A) (Li et al. 2016; Dominissini et al. 2016), initially identified in total RNA and rRNAs (Dunn 1961; Hall 1963; Klootwijk and Planta 1973). Intriguingly, the position of m^1^A modifications has been shown to be more correlated with the position of the first intron than with transcriptional or translational start sites (Figure 2g from Dominissini et al. 2016). When the distance of m^1^As to each splice site in a given mRNA was calculated, the first splice site was found to be the nearest for 85% of m^1^ As (Dominissini et al. 2016). When 5’UTR introns were present, m^1^A was found to be near the first splice site regardless of the position of the start codon (Dominissini et al. 2016). Given that 5IM transcripts are also characterized by the position of the first intron, we investigated the relationship between 5IMP score and m^1^A RNA modification marks.

We analyzed the union of previously identified m^1^A modifications (Dominissini et al. 2016) across all cell types and conditions. Although there is some evidence that these marks depend on cell type and growth condition, it is difficult to be confident of the cell type and condition-dependence of any particular mark given experimental variation (see Methods). Nevertheless, we found that mRNAs with m^1^A modification early in the coding region (first 99 nucleotides) had substantially higher 5IMP scores than mRNAs lacking these marks (**Fig 6A**; Wilcoxon Rank Sum Test p = 3.4e-265), and were greatly enriched among 5IM transcripts (**Fig 6A**; Fisher’s Exact Test p = 1.6e-177; odds ratio = 3.8). In other words, the sequence features within the early coding region that define 5IMP transcripts also associate with m^1^A modification in the early coding region.

**Fig 6.**
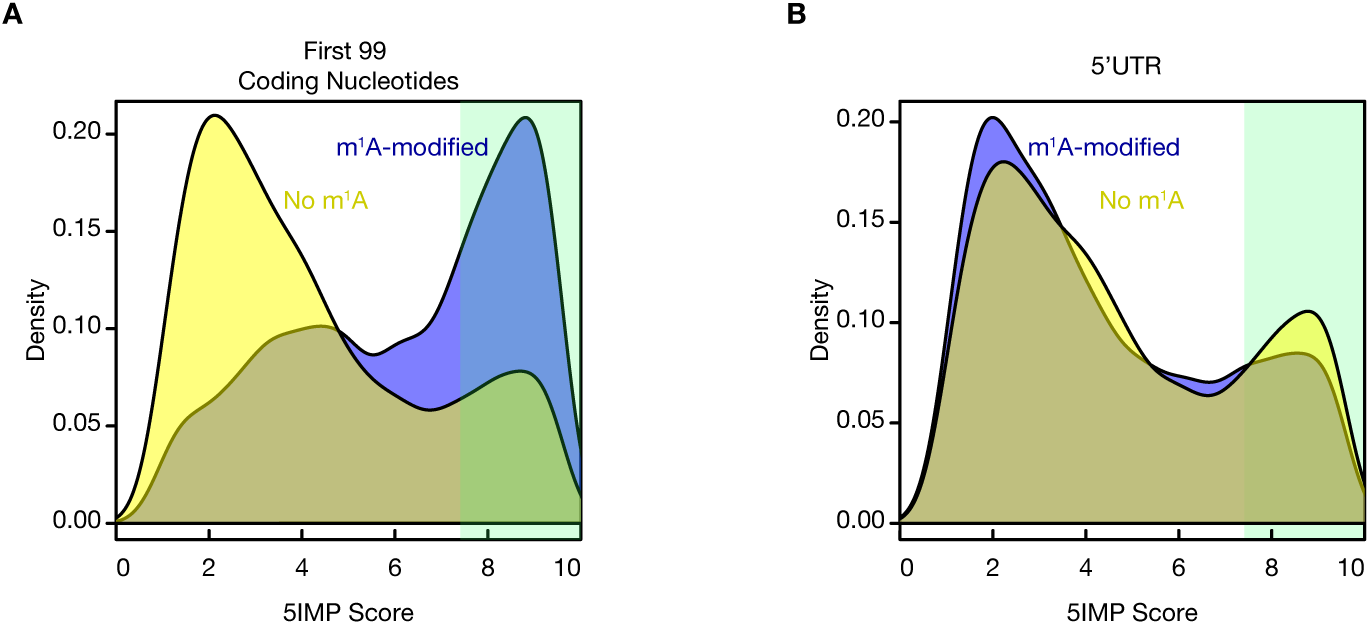
5IM transcripts are enriched for mRNAs with early coding region m^1^A modifications. **(a)** Transcripts with m^1^A modifications (blue) in the first 99 coding nucleotides exhibit significant enrichment for 5IM transcripts and have higher 5IMPS scores than transcripts without m^1^A modifications in the first 99 coding nucleotides (yellow). **(b)** Transcripts with m^1^A modifications (blue) in the 5’UTR do not display a similar enrichment.

We next wondered whether 5IMP score was related to m^1^A modification generally, or only associated with m^1^A modification in the early coding region. Indeed, many of the previously identified m^1^A peaks were within the 5’UTRs of mRNAs (Li et al. 2016). Interestingly, 5IMP scores were only associated with m^1^A modification in the early coding region, and not with m^1^A modification in the 5’UTR (**Fig 6B**). This offers the intriguing possibility that the sequence features that define 5IMP transcripts are co-localized with m^1^A modification.

## Discussion

Coordinating the expression of functionally related transcripts can be achieved by post-transcriptional processes such as splicing, RNA export, RNA localization or translation (Moore and Proudfoot 2009). Sets of mRNAs subject to a common regulatory transcriptional process can exhibit common sequence features that define them to be a class. For example, transcripts subject to regulation by particular miRNAs tend to share certain sequences in their 3’UTRs that are complementary to the regulatory miRNA (Ameres and Zamore 2013). Similarly, transcripts that share a 5’ terminal oligopyrimidine tract are coordinately regulated by mTOR and ribosomal protein S6 kinase (Meyuhas 2000). Here we quantitatively define ‘5IM’ transcripts as a class that shares common sequence elements and functional properties. We estimate the 5IM class to comprise 21% of all human transcripts.

Whereas 35% of human transcripts have one or more 5’UTR introns, the majority of 5IM transcripts have neither a 5’UTR intron nor an intron in the first 90 nts of the ORF. Other shared features of 5IM transcripts include sequence features associated with low translation initiation rates. These are: (1) a tendency for RNA secondary structure in the region immediately preceding the start codon (**Fig 3A**), and near the 5’cap (**Fig 3B**); (2) translational upregulation upon overexpression of eIF4E (**Fig 3C**); and (3) more frequent use of non-AUG start codons (**Fig 3D**). Also consistent with low intrinsic translation efficiencies, 5IM transcripts additionally tend to depend on less abundant tRNAs to decode the beginning of the open reading frame (**Fig 3E**).

We had previously reported that transcripts encoding proteins with ER- and mitochondrial-targeting signal sequences (SSCRs and MSCRs, respectively) are over-represented among the 65% of transcripts lacking 5’UTR introns (Cenik et al. 2011). Transcripts in this set are enriched for the sequence features detected by our 5IM classifier. By examining these enriched sequence features, we showed that the 5IM class extends beyond mRNAs encoding membrane proteins. Jan et al. recently developed a transcriptome-scale method to identify mRNAs occupied by ER-proximal ribosomes in both yeast and human cells (Jan et al. 2014). As expected, transcripts known to encode ER-trafficked proteins were highly enriched for ER-proximal ribosome occupancy. However, their data also showed many transcripts encoding non-ER trafficked proteins to also be engaged with ER-proximal ribosomes (Reid and Nicchitta 2015b). Similarly, several other studies have suggested a critical role of ER-proximal ribosomes in translating several cytoplasmic proteins (Reid and Nicchitta 2015a). Here, we found that 5IM transcripts including those that are not ER-trafficked or mitochondrial were significantly more likely to exhibit binding to ER-proximal ribosomes **(Figs 4A-B**).

In addition to ribosomes directly resident on the ER, an interesting possibility is the presence of a pool of peri-ER ribosomes (Jan et al. 2015; Reid and Nicchitta 2015a). Association of 5IM transcripts with such a peri-ER ribosome pool could potentially explain the observed correlation of 5IM status with binding to ER-proximal ribosomes. The ER is physically proximal to mitochondria (Rowland and Voeltz 2012), so peri-ER ribosomes may include those translating mRNAs on mitochondria (i.e., mRNAs with MSCRs) (Sylvestre et al. 2003). However, even when transcripts corresponding to ER-trafficked and mitochondrial proteins were excluded from consideration, ER-proximal ribosome enrichment and 5IMP scores were highly correlated (**Fig 4A**). Thus another shared feature of 5IM transcripts is their translation on or near the ER regardless of the ultimate destination of the encoded protein.

In an attempt to identify a common factor binding 5IM transcripts, we asked whether 5IM transcripts were enriched among the experimentally identified targets of 23 different RBPs. Only one RBP emerged--the exon junction complex (EJC). Specifically, we observed a dramatic enrichment of non-canonical EJC (ncEJC) binding sites within the early coding region of 5IM transcripts. Further, the CG-rich motif identified for ncEJCs in first exons is strikingly similar to the CG-rich motif enriched in the first exons of 5IM transcripts (**Fig 5B**). Previous work implicated RanBP2, a protein associated with the cytoplasmic face of the nuclear pore, as a binding factor for some SSCRs (Mahadevan et al. 2013). This finding suggests that nuclear pore proteins may influence EJC occupancy on these transcripts.

EJC deposition during the process of pre-mRNA splicing enables the nuclear history of an mRNA to influence post-transcriptional processes including mRNA localization, translation efficiency, and nonsense mediated decay (Chang et al. 2007; Kervestin and Jacobson 2012; Choe et al. 2014). While canonical EJC binding occurs at a fixed distance upstream of exon-exon junctions and involves direct contact between the sugar-phosphate backbone and the EJC core anchoring protein eIF4AIII, ncEJC binding sites likely reflect stable engagement between the EJC core and other mRNP proteins (e.g., SR proteins) recognizing nearby sequence motifs. Although some RBPs were identified for ncEJC motifs found in internal exons (Singh et al. 2014, 2012), to date no candidate RBP has been identified for the CG-rich ncEJC motif found in the first exon. If this motif does result from an RBP interaction, it is likely to be one or more of the ∼70 proteins that stably and specifically bind to the EJC core (Singh et al. 2012).

Finally, we observed a dramatic enrichment for m^1^A modifications among 5IM transcripts, with specific enrichment for m^1^A modifications in the early coding region. Given this striking enrichment perhaps it is perhaps not surprising that m^1^A containing mRNAs were also shown to have more structured 5’UTRs that are GC-rich compared to m^1^A lacking mRNAs (Dominissini et al. 2016). Similar to 5IM transcripts, m^1^A containing mRNAs were found to decorate start codons that appear in a highly structured context. While ALKBH3 has been identified as a protein that can demethylate m^1^A, it is currently unknown whether there are any proteins that can specifically act as “readers” of m^1^A. Recent studies have begun to identify such readers for other mRNA modifications such as YTHDF1, YTHDF2, WTAP and HNRNPA2B1 (Ping et al. 2014; Liu et al. 2014; Wang et al. 2014, 2015; Alarcón et al. 2015). Our study highlights a possible link between non-canonical EJC binding and m^1^A. Hence, our results yield the intriguing hypothesis that one or more of the ∼70 proteins that stably and specifically bind to the EJC core can function as an m^1^A reader. Future work involving directed experiments would be needed to test this hypothesis.

Given that 5IM transcripts are enriched for ER-targeted and mitochondrial proteins, it is plausible that the observed functional characteristics of 5IM transcripts are driven solely by SSCR and MSCR-containing transcripts. Hence, we repeated all analyses for the subclasses of 5IM transcripts (MSCR-containing, SSCR-containing, S^−^/M^−^/SignalP^+^, or S^−^/M^−^/SignalP^−^). We found the observed associations remained statistically significant and had the same direction of effect, even after eliminating SSCR- and MSCR-containing transcripts, despite the fact that all training of the 5IM classifier was performed only using SSCR transcripts. We also found that 5IMP score was equally or more strongly associated with each of the functional characteristics compared to the 5UI status. In conclusion, the molecular associations we report apply to 5IM transcripts as a whole, and are not driven solely by the subset of 5IM transcripts encoding ER- or mitochondria-targeting signal peptides, and seem to indicate shared features beyond simple lack of a 5’UTR intron.

An intriguing possibility is that 5IM transcript features associated with lower intrinsic translation efficiency may together enable greater ‘tunability’ of 5IM transcripts at the translation stage. Regulated enhancement or repression of translation, for 5IM transcripts, could allow for rapid changes in protein levels. There are analogies to this scenario in transcriptional regulation, wherein highly regulated genes often have promoters with low baseline levels that can be rapidly modulated through the action of regulatory transcription factors. As more ribosome profiling studies are published examining translational responses transcriptome-wide under multiple perturbations, conditions under which 5IM transcripts are translationally regulated may be revealed. Directed experiments will be needed to test translational features of 5IM transcripts hypothesized via this computational analysis.

Taken together, our analyses reveal the existence of a distinct ‘5IM’ class comprising 21% of human transcripts. This class is defined by depletion of 5’ proximal introns, presence of specific RNA sequence features associated with low translation efficiency, non-canonical binding by the Exon Junction Complex and an enrichment for N^1^-methyladenosine modification.

## Materials and Methods

### Datasets and Annotations

Human transcript sequences were downloaded from the NCBI human Reference Gene Collection (RefSeq) via the UCSC table browser (hg19) on Jun 25 2010 (Kent et al. 2002; Pruitt et al. 2005). Transcripts with fewer than three coding exons, and transcripts where the first 99 coding nucleotides straddle more than two exons were removed from further consideration. The criteria for exclusion of genes with fewer than three coding exons was to ensure that the analysis of downstream regions was possible for all genes that were used in our analysis of early coding regions. Specifically, the downstream regions were selected randomly from downstream of the 3^rd^ exons. Hence, genes with fewer exons would not be able to contribute a downstream region potentially creating a skew in representation. In total there were ∼3000 genes that were removed from consideration due to this filter. Therefore, our classifier is limited in its ability to assess transcripts from these genes. However, the performance measures reported in our manuscript are robust to exclusion of these genes, in the sense that the same class of transcripts was used in both training and test datasets.

Transcripts were clustered based on sequence similarity in the first 99 coding nucleotides. Specifically, each transcript pair was aligned using BLAST with the DUST filter disabled (Altschul et al. 1990). Transcript pairs with BLAST E-values < 1e-25 were grouped into transcript clusters. In total, there were 15576 transcript clusters that were considered further. These clusters that were subsequently assigned to one of four categories: MSCR containing, SSCR-containing, S^−^/MSCR^−^SignalP^+^, or S^−^/MSCR^−^ SignalP^−^ as follows:

MSCR-containing transcripts were annotated using MitoCarta and other sources as described in (Cenik et al. 2011). SSCR-containing transcripts were the set of transcripts annotated to contain signal peptides in the Ensembl Gene v.58 annotations, which were downloaded through Biomart on Jun 25 2010. For transcripts without an annotated MSCR or SSCR, the first 70 amino acids were analyzed using SignalP 3.0 (Bendtsen et al. 2004). Using the eukaryotic prediction mode, transcripts were assigned to the S^−^/MSCR^−^ SignalP^+^ category if either the Hidden Markov Model or the Artificial Neural Network classified the sequence as signal peptide containing. All remaining transcript clusters were assigned to the S^−^/MSCR^−^ SignalP^−^ category. The number of transcript clusters in each of the four categories was: 3743 SSCR, 737 MSCR, 696 S^−^/MSCR^−^ SignalP^+^, 10400 S^−^/MSCR^−^ SignalP^−^.

For each transcript cluster, we also constructed matched control sequences. Control sequences were derived from a single randomly chosen in-frame ‘window’ downstream of the 3rd exon from the evaluated transcripts. If an evaluated transcript had fewer than 99 nucleotides downstream of the 3^rd^ exon, no control sequence was extracted. 5UI labels and transcript clustering for the control sequences were inherited from the evaluated transcript. The rationale for this decision is that our analysis depends on the position of the first intron; hence genes with fewer than two exons need to be excluded, as these will not have introns. We further required the matched control sequences to fall downstream of the early coding region. In the vast majority of cases the third exon fell outside the first 99 nucleotides of the coding region, making this a convenient criterion by which to choose control regions.

### Sequence Features and Motif Discovery

36 sequence features were extracted from each transcript (**Table S1**). The sequence features included the ratio of arginines to lysines, the ratio of leucines to isoleucines, adenine content, length of the longest stretch without adenines, preference against codons that contain adenines or thymines. These features were previously found to be enriched in SSCR-containing and certain 5UI^−^ transcripts (Cenik et al. 2011; Palazzo et al. 2007). In addition, we extracted ratios between several other amino acid pairs based on having biochemical/evolutionary similarity, i.e. having positive scores, according to the BLOSUM62 matrix (Henikoff and Henikoff 1992). To avoid extreme ratios given the relatively short sequence length, pseudo-counts were added to amino acid ratios using their respective genome-wide prevalence.

In addition, we used three published motif finding algorithms (AlignACE, DEME, and MoAN (Roth et al. 1998; Redhead and Bailey 2007; Valen et al. 2009)) to discover RNA sequence motifs enriched among 5UI^−^ transcripts. AlignACE implements a Gibbs sampling approach and is one of the pioneering efforts in motif discovery (Roth et al. 1998). We modified the AlignACE source code to restrict motif searches to only the forward strand of the input sequences to enable RNA motif discovery. DEME and MoAN adopt discriminative approaches to motif finding by searching for motifs that are differentially enriched between two sets of sequences (Redhead and Bailey 2007; Valen et al. 2009). MoAN has the additional advantage of discovering variable length motifs, and can identify co-occurring motifs with the highest discriminative power.

In total, six motifs were discovered using the three motif finding algorithms (**Table S1**). Position specific scoring matrices for all motifs were used to score the first 99 - *l* positions in each sequence, where *l* is the length of the motif. We assessed the significance of each motif instance by calculating the p-value of enrichment (Fisher’s Exact Test) among 5UI^−^ transcripts considering all transcripts with a motif instance achieving a PSSM score greater than equal to the instance under consideration. The significance score and position of the two best motif instances were used as features for the classifier (**Table S1**).

### 5IM Classifier Training and Performance Evaluation

We modified an implementation of the Random Forest classifier (Breiman 2001) to model the relationship between sequence features in the early coding region and the absence of 5’UTR introns (5UIs). This classifier discriminates transcripts with <u>5</u>’proximal-<u>i</u>ntron-<u>m</u>inus-like-coding regions and hence is named the ‘5IM’ classifier. The training set for the classifier was composed of SSCR transcripts exclusively. There were two reasons to restrict model construction to SSCR transcripts: 1) we expected the presence of specific RNA elements as a function of 5UI presence based on our previous work (Cenik et al. 2011); and 2) we wanted to restrict model building to sequences that have similar functional constraints at the protein level.

We observed that 5UI^−^ transcripts with introns proximal to 5’ end of the coding region have sequence characteristics similar to 5UI^+^ transcripts (**Fig 1D**). To systematically characterize this relationship, we built different classifiers using training sets that excluded 5UI^−^ transcripts with a coding region intron positioned at increasing distances from the start codon. We evaluated the performance of each classifier using 10-fold cross validation.

Given that a large number of motif discovery iterations were needed, we sought to reduce the computational burden. We isolated a subset of the training examples to be used exclusively for motif finding. Motif discovery was performed once using this set of sequences, and the same motifs were used in each fold of the cross validation for all the classifiers. Imbalances between the sizes of positive and negative training examples can lead to detrimental classification performance (Wang and Yao 2012). Hence, we balanced the training set size of 5UI^−^ and 5UI^+^ transcripts by randomly sampling from the larger class. We constructed 10 sub-classifiers to reduce sampling bias, and for each test example, the prediction score from each subclassifier was summed to produce a combined score between 0 and 10. For the rest of the analyses, we used the classifier trained using 5UI^−^ transcripts where first coding intron falls outside the first 90 coding nucleotides (**Fig 1E**).

We evaluated classifier performance using a 10-fold cross validation strategy for SSCR-containing transcripts (i.e. the training set). In each fold of the cross-validation, the model was trained without any information from the held-out examples, including motif discovery. For all the other transcripts and the control sets (see above), the 5IMP scores were calculated using the classifier trained using SSCR transcripts as described above. 5IMP score distribution for the control set was used to calculate the empirical cumulative null distribution. Using this function, we determined the p-value corresponding to the 5IMP score for all transcripts. We corrected for multiple hypotheses testing using the qvalue R package (Storey 2003). Based on this analysis, we estimate that a 5IMP score of 7.41 corresponds to a 5% False Discovery Rate and suggest that 21% of all human transcripts can be considered as 5IM transcripts.

While the theoretical range of 5IMP scores is 0-10, the highest observed 5IMP is 9.855. We note that for all figures that depict 5IMP score distributions, we displayed the entire theoretical range of 5IMP scores (0-10).

### Functional Characterization of 5IM Transcripts

We collected genome-scale dataset from publically available databases and from supplementary information provided in selected articles. For all analyzed datasets, we first converted all gene/transcript identifiers (IDs) into RefSeq transcript IDs using the Synergizer webserver (Berriz and Roth 2008). If a dataset was generated using a non-human species (ex. Targets identified by TDP-43 RNA immunoprecipitation in rat neuronal cells), we used Homologene release 64 (downloaded on Sep 28 2009) to identify the corresponding ortholog in humans. If at least one member of a transcript cluster was associated with a functional phenotype, we assigned the cluster to the positive set with respect to the functional phenotype. If more than one member of a cluster had the functional phenotype, we only retained one copy of the cluster unless they differed in a quantitative measurement. For example, consider two hypothetical transcripts: NM_1 and NM_2 that were clustered together and have a 5IMP score of 8.5. If NM_1 had an mRNA half-life of 2hr while NM_2’s half-life was 1hr than we split the cluster while preserving the 5IMP score for both NM_1 and NM_2.

Once the transcripts were partitioned based on the functional phenotype, we ran two statistical tests: 1) Fisher’s Exact Test for enrichment of 5IM transcripts within the functional category; 2) Wilcoxon Rank Sum Test to compare 5IMP scores between transcripts partitioned by the functional phenotype. Additionally, for datasets where a quantitative measurement was available (ex. mRNA half-life), we calculated the Spearman rank correlation between 5IMP scores and the quantitative variable. In these analyses, we assumed that the test space was the entire set of RefSeq transcripts. For all phenotypes where we observed a preliminary statistically significant result, we followed up with more detailed analyses described below.

### Analysis of Features Associated with Translation

For each transcript, we predicted the propensity for secondary structure preceding the translation start site and the 5’cap. Specifically, we extracted 35 nucleotides preceding the translation start site or the first 35 nucleotides of the 5’UTR. If a 5’UTR is shorter than 35 nucleotides, the transcript was removed from the analysis. hybrid-ss-min utility (UNAFold package version 3.8) with default parameters was used to calculate the minimum folding energy (Markham and Zuker 2008).

Codon optimality was measured using the tRNA Adaptation Index (tAI), which is based on the genomic copy number of each tRNA (dos Reis et al. 2004). tAI for all human codons were downloaded from Tuller et al. 2010 Table S1. tAI profiles for the first 30 amino acids were calculated for all transcripts. Codon optimality profiles were summarized for the first 30 amino acids for each transcript or by averaging tAI at each codon.

We carried out two control experiments to test whether the association between 5IMP score and tAI could be explained by confounding variables. First, we permuted the nucleotides in the first 90 nts and observed no relationship between 5IMP score and mean tAI when these permuted sequences were used (**Fig S1**). Second, we selected random in-frame 99 nucleotides from 3^rd^ exon to the end of the coding region and found no significant differences in tAI (**Fig S1**). These results suggest that the relationship between tAI and 5IMP score is confined to the first 30 amino acids and is not explained by simple differences in nucleotide composition.

Ribosome profiling and RNA expression data for human lymphoblastoid cells (LCLs) were downloaded from NCBI GEO database accession number: GSE65912. Translation efficiency was calculated as previously described (Cenik et al. 2015). Median translation efficiency across the different cell-types was used for each transcript.

### Analysis of Proximity Specific Ribosome Profiling Data

We downloaded proximity specific ribosome profiling data for HEK 293 cells from Jan et al. 2014; Table S6. We converted UCSC gene identifiers to HGNC symbols using g:Profiler (Reimand et al. 2011). We retained all genes with an RPKM >5 in either input or pulldown and required that at least 30 reads were mapped in either of the two libraries. We used ER-targeting evidence categories “secretome”, “phobius”, “TMHMM”, “SignalP”, “signalSequence”, “signalAnchor” from (Jan et al. 2014) to annotate genes as having ER-targeting evidence. The genes that did not have any ER-targeting evidence or “mitoCarta” /“mito.GO” annotations were deemed as the set of genes with no ER-targeting or mitochondrial evidence. We calculated the log_2_ of the ratio between ER-proximal ribosome pulldown RPKM and control RPKM as the measure of enrichment for ER-proximal ribosome occupancy (as in Jan et al. 2014). A moving average of this ratio was calculated for genes grouped by their 5IMP score. For this calculation, we used bins of 30 mitochondrial genes or 100 genes with no evidence of ER- or mitochondrial targeting.

### Analysis of Genome-wide Binding Sites of Exon Junction Complex

Dr. Gene Yeo and Gabriel Pratt generously shared uniformly processed peak calls for experiments identifying human RNA binding protein targets. These datasets include various CLIP-Seq datasets and its variants such as iCLIP. A total 49 datasets from 22 factors were analyzed. These factors were: hnRNPA1, hnRNPF, hnRNPM, hnRNPU, Ago2, hnRNPU, HuR, IGF2BP1, IGF2BP2, IGF2BP3, FMR1, eIF4AIII, PTB, IGF2BP1, Ago3, Ago4, MOV10, Fip1, CF Im68, CF Im59, CF Im25, and hnRNPA2B1. We extracted the 5IMP scores for all targets of each RBP. We calculated the Wilcoxon Rank Sum test statistic comparing the 5IMP score distribution of the targets of each RBP to all other transcripts with 5IMP scores. None of the tested RBP target sets had an adjusted p-value < 0.05 and a median difference in 5IMP score > 1 when compared to non-target transcripts.

In addition, we used RNA:protein immunoprecipitation in tandem (RIPiT) data to determine Exon Junction Complex (EJC) binding sites (Singh et al. 2014, 2012). We analyzed the common peaks from the Y14-Magoh immunoprecipitation configuration (Singh et al. 2012; Kucukural et al. 2013). Canonical EJC binding sites were defined as peaks whose weighted center were 15 to 32 nucleotides upstream of an exon-intron boundary. All remaining peaks were deemed as “non-canonical” EJC binding sites. We extracted all non-canonical peaks that overlapped the first 99 nucleotides of the coding region and restricted our analysis to transcripts that had an RPKM greater than one in the matched RNA-Seq data.

### Analysis of m^1^A modified transcripts

We downloaded the list of RNAs observed to contain m^1^A from Li et al. (2016) and Dominissini et al. (2016). RefSeq transcript identifiers were converted to HGNC symbols using g:Profiler (Reimand et al. 2011). The overlap between the two datasets was determined using HGNC symbols. m^1^A modifications that overlap the first 99 nucleotides of the coding region were determined using bedtools (Quinlan and Hall 2010).

Li et al. (2016) identified 600 transcripts with m^1^A modification in normal HEK293 cells. Of these, 368 transcripts were not found to contain the m^1^A modification in HEK293 cells by Dominissini et al. (2016). Yet, 81% of these were found to be m^1^A modified in other cell types. Li et al. (2016) also analyzed m^1^A upon H_2_O_2_ treatment and serum starvation in HEK293 cells and identified many m^1^A modifications that are only found these stress conditions. However, 20% of 371 transcripts harboring stress-induced m^1^A modifications were found in normal HEK293 cells by Dominissini et al. (2016). Taken together, these analyses suggest that transcriptome-wide m^1^A maps remain incomplete. Hence, we analyzed the 5IMP scores of all mRNAs with m^1^A across cell types and conditions. We reported results using the Dominissini et al. dataset but the same conclusions were supported by m^1^A modifications from Li et al.

## Acknowledgements

We would like to thank Gene Yeo, and Gabriel Pratt for sharing peak calls for RNA binding protein target identification experiments (CLIP, PAR-CLIP, iCLIP). We thank Elif Sarinay Cenik, Başar Cenik, and Alper Küçükural for helpful discussions. F.P.R. and H.N.C. were supported by the Canada Excellence Research Chairs Program. This work was supported in part by NIH grants GM50037 (M.J.M.), HG001715 and HG004233 (F.P.R.), the Krembil Foundation, an Ontario Research Fund award, and the Avon Foundation (F.P.R.). A.F.P. was supported by a grant from the Canadian Institutes of Health Research to A.F.P. (FRN 102725).

## Author contributions

CC, MJM, and FPR designed the study. CC and HNC implemented the random forest classifier and carried out all analyses. GS and AA performed initial experiments. MPS and AFP provided critical feedback. MJM and FPR jointly supervised the project. CC, FPR and MJM wrote the manuscript with input from all authors.

## Supporting Information

**Figure S1.**
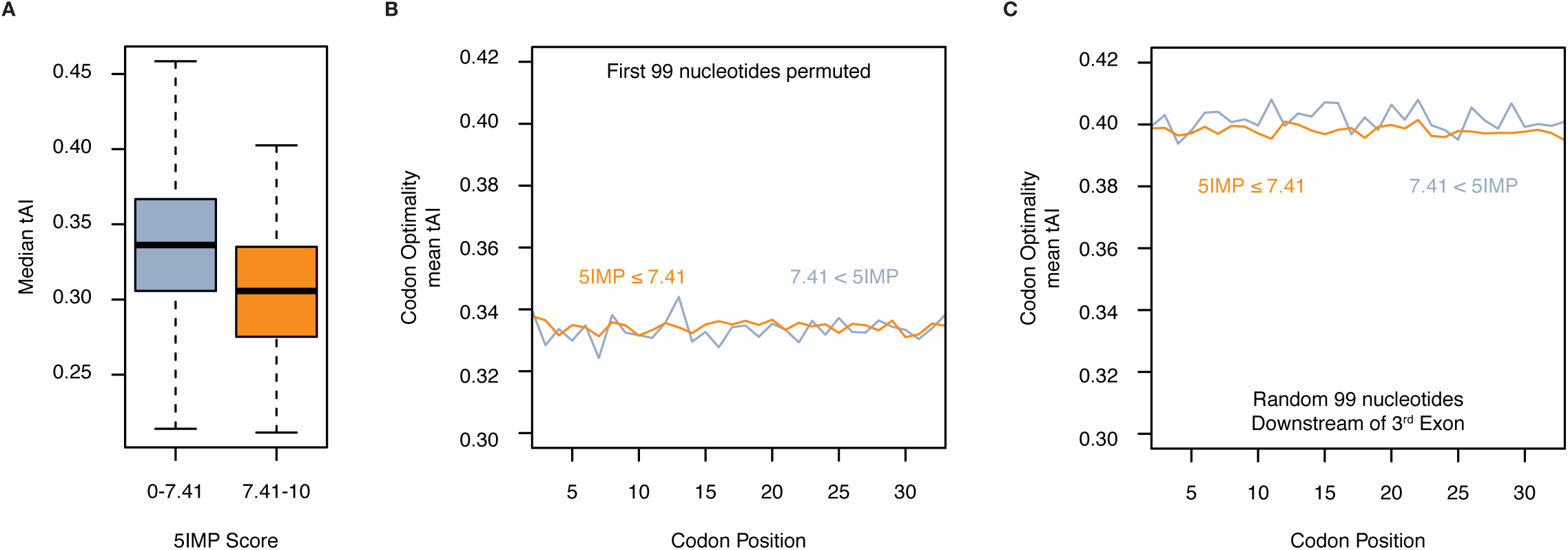
Association between 5IMP scores and codon optimality is restricted to the first 30 amino acids and is not explained by nucleotide content. **(a)** 5IM transcripts tend to have less optimal codons in their first 30 amino acids as measured by tRNA adaptation index (tAI). The median tAI for each transcript was calculated and transcripts were grouped by their 5IMP scores. The distribution of median tAI was plotted as a boxplot. **(b)** We permuted the nucleotides of the first 99 nucleotides and found that the relationship between 5IMP and tAI was lost. This result suggested that nucleotide composition alone doesn’t explain the relationship between 5IMP and tAI. **(c)** We used the previously described negative control sequences, each derived from a single randomly chosen in-frame ‘window’ downstream of the 3rd exon from one of the evaluated transcripts. We found no relationship between 5IMP score and tAI in these regions suggesting that the observed association is restricted to the early coding region.

**Table S1-** | List of features used by the 5IM classifier.

**Table S2-** | 5IMP scores of all human transcripts.

**Table S3-** | List of functional features tested for association with 5IMP scores.

## References

Alarcón CR, Goodarzi H, Lee H, Liu X, Tavazoie S, Tavazoie SF. 2015. HNRNPA2B1 Is a Mediator of m(6)A-Dependent Nuclear RNA Processing Events. Cell 162: 1299–1308.

Altschul SF, Gish W, Miller W, Myers EW, Lipman DJ. 1990. Basic local alignment search tool. J Mol Biol 215: 403–410.

Ameres SL, Zamore PD. 2013. Diversifying microRNA sequence and function. Nat Rev Mol Cell Biol 14: 475–488.

Babendure JR, Babendure JL, Ding J-H, Tsien RY. 2006. Control of mammalian translation by mRNA structure near caps. RNA 12: 851–861.

Bendtsen JD, Nielsen H, von Heijne G, Brunak S. 2004. Improved prediction of signal peptides: SignalP 3.0. J Mol Biol 340: 783–795.

Berriz GF, Roth FP. 2008. The Synergizer service for translating gene, protein and other biological identifiers. Bioinformatics 24: 2272–2273.

Breiman L 2001. Random Forests. Mach Learn 45: 5–32.

Cenik C, Chua HN, Zhang H, Tarnawsky SP, Akef A, Derti A, Tasan M, Moore MJ, Palazzo AF, Roth FP. 2011. Genome analysis reveals interplay between 5’UTR introns and nuclear mRNA export for secretory and mitochondrial genes. PLoS Genet 7: e1001366.

Cenik C, Derti A, Mellor JC, Berriz GF, Roth FP. 2010. Genome-wide functional analysis of human 5’ untranslated region introns. Genome Biol 11: R29.

Cenik C, Sarinay Cenik E, Byeon GW, Grubert F, Candille SI, Spacek D, Alsallakh B, Tilgner H, Araya CL, Tang H, et al. 2015. Integrative analysis of RNA, translation and protein levels reveals distinct regulatory variation across humans. Genome Res. http://dx.doi.org/10.1101/gr.193342.115.

Chang Y-F, Imam JS, Wilkinson MF. 2007. The nonsense-mediated decay RNA surveillance pathway. Annu Rev Biochem 76 : 51–74.

Charneski CA, Hurst LD. 2013. Positively charged residues are the major determinants of ribosomal velocity. PLoS Biol 11: e1001508.

Choe J, Ryu I, Park OH, Park J, Cho H, Yoo JS, Chi SW, Kim MK, Song HK, Kim YK. 2014. eIF4AIII enhances translation of nuclear cap-binding complex-bound mRNAs by promoting disruption of secondary structures in 5’UTR. Proc Natl Acad Sci U S A 111: E4577–86.

Dominissini D, Nachtergaele S, Moshitch-Moshkovitz S, Peer E, Kol N, Ben-Haim MS, Dai Q, Di Segni A, Salmon-Divon M, Clark WC, et al. 2016. The dynamic N(1)-methyladenosine methylome in eukaryotic messenger RNA. Nature 530: 441–446.

dos Reis M, Savva R, Wernisch L. 2004. Solving the riddle of codon usage preferences: a test for translational selection. Nucleic Acids Res 32: 5036–5044.

Dunn DB. 1961. The occurrence of 1-methyladenine in ribonucleic acid. Biochim Biophys Acta 46: 198–200.

Frye M, Jaffrey SR, Pan T, Rechavi G, Suzuki T. 2016. RNA modifications: what have we learned and where are we headed? Nat Rev Genet 17: 365–372.

Gerashchenko MV, Gladyshev VN. 2015. Translation inhibitors cause abnormalities in ribosome profiling experiments. Nucleic Acids Res 42: e134.

Gingold H, Pilpel Y. 2011. Determinants of translation efficiency and accuracy. Mol Syst Biol 7: 481.

Hall RH. 1963. Method for isolation of 2′-O-methylribonucleosides and N1-methyladenosine from ribonucleic acid. Biochimica et Biophysica Acta (BBA) - Specialized Section on Nucleic Acids and Related Subjects 68 : 278–283.

Henikoff S, Henikoff JG. 1992. Amino acid substitution matrices from protein blocks. Proc Natl Acad Sci U S A 89 : 10915–10919.

Hershberg R, Petrov DA. 2008. Selection on codon bias. Annu Rev Genet 42: 287–299.

Hinnebusch AG, Lorsch JR. 2012. The mechanism of eukaryotic translation initiation: new insights and challenges. Cold Spring Harb Perspect Biol 4. http://dx.doi.org/10.1101/cshperspect.a011544.

Hong X, Scofield DG, Lynch M. 2006. Intron size, abundance, and distribution within untranslated regions of genes. Mol Biol Evol 23: 2392–2404.

Jan CH, Williams CC, Weissman JS. 2015. LOCAL TRANSLATION. Response to Comment on “Principles of ER cotranslational translocation revealed by proximity-specific ribosome profiling.” Science 348: 1217.

Jan CH, Williams CC, Weissman JS. 2014. Principles of ER cotranslational translocation revealed by proximity-specific ribosome profiling. Science 346: 1257521.

Kent WJ, Sugnet CW, Furey TS, Roskin KM, Pringle TH, Zahler AM, Haussler D. 2002. The human genome browser at UCSC. Genome Res 12: 996–1006.

Kervestin S, Jacobson A. 2012. NMD: a multifaceted response to premature translational termination. Nat Rev Mol Cell Biol 13 : 700–712.

Klootwijk J, Planta RJ. 1973. Analysis of the methylation sites in yeast ribosomal RNA. Eur J Biochem 39: 325–333.

Kucukural A, Özadam H, Singh G, Moore MJ, Cenik C. 2013. ASPeak: an abundance sensitive peak detection algorithm for RIP-Seq. Bioinformatics 29: 2485–2486.

Larsson O, Li S, Issaenko OA, Avdulov S, Peterson M, Smith K, Bitterman PB, Polunovsky VA. 2007. Eukaryotic Translation Initiation Factor 4E–Induced Progression of Primary Human Mammary Epithelial Cells along the Cancer Pathway Is Associated with Targeted Translational Deregulation of Oncogenic Drivers and Inhibitors. Cancer Res 67: 6814–6824.

Lee ES, Akef A, Mahadevan K, Palazzo AF. 2015. The consensus 5’ splice site motif inhibits mRNA nuclear export. PLoS One 10: e0122743.

Le Hir H, Izaurralde E, Maquat LE, Moore MJ. 2000. The spliceosome deposits multiple proteins 20–24 nucleotides upstream of mRNA exon-exon junctions. EMBO J 19: 6860–6869.

Liu J, Yue Y, Han D, Wang X, Fu Y, Zhang L, Jia G, Yu M, Lu Z, Deng X, et al. 2014. A METTL3-METTL14 complex mediates mammalian nuclear RNA N6-adenosine methylation. Nat Chem Biol 10: 93–95.

Li X, Xiong X, Wang K, Wang L, Shu X, Ma S, Yi C. 2016. Transcriptome-wide mapping reveals reversible and dynamic N1-methyladenosine methylome. Nat Chem Biol. http://dx.doi.org/10.1038/nchembio.2040 (Accessed March 29, 2016).

Mahadevan K, Zhang H, Akef A, Cui XA, Gueroussov S, Cenik C, Roth FP, Palazzo AF. 2013. RanBP2/Nup358 potentiates the translation of a subset of mRNAs encoding secretory proteins. PLoS Biol 11: e1001545.

Marintchev A, Edmonds KA, Marintcheva B, Hendrickson E, Oberer M, Suzuki C, Herdy B, Sonenberg N, Wagner G. 2009. Topology and regulation of the human eIF4A/4G/4H helicase complex in translation initiation. Cell 136: 447–460.

Markham NR, Zuker M 2008. UNAFold: software for nucleic acid folding and hybridization. Methods Mol Biol 453: 3–31.

Meyuhas O 2000. Synthesis of the translational apparatus is regulated at the translational level. Eur J Biochem 267: 6321–6330.

Moore MJ, Proudfoot NJ. 2009. Pre-mRNA processing reaches back to transcription and ahead to translation. Cell 136: 688–700.

Palazzo AF, Mahadevan K, Tarnawsky SP. 2013. ALREX-elements and introns: two identity elements that promote mRNA nuclear export. Wiley Interdiscip Rev RNA 4: 523–533.

Palazzo AF, Springer M, Shibata Y, Lee C-S, Dias AP, Rapoport TA. 2007. The signal sequence coding region promotes nuclear export of mRNA. PLoS Biol 5: e322.

Parsyan A, Svitkin Y, Shahbazian D, Gkogkas C, Lasko P, Merrick WC, Sonenberg N. 2011. mRNA helicases: the tacticians of translational control. Nat Rev Mol Cell Biol 12: 235–245.

Ping X-L, Sun B-F, Wang L, Xiao W, Yang X, Wang W-J, Adhikari S, Shi Y, Lv Y, Chen Y-S, et al. 2014. Mammalian WTAP is a regulatory subunit of the RNA N6-methyladenosine methyltransferase. Cell Res 24: 177–189.

Pruitt KD, Tatusova T, Maglott DR. 2005. NCBI Reference Sequence (RefSeq): a curated non-redundant sequence database of genomes, transcripts and proteins. Nucleic Acids Res 33: D501–4.

Quinlan AR, Hall IM. 2010. BEDTools: a flexible suite of utilities for comparing genomic features. Bioinformatics 26: 841–842.

Redhead E, Bailey TL. 2007. Discriminative motif discovery in DNA and protein sequences using the DEME algorithm. BMC Bioinformatics 8: 385.

Reid DW, Nicchitta CV. 2015a. Diversity and selectivity in mRNA translation on the endoplasmic reticulum. Nat Rev Mol Cell Biol 16: 221–231.

Reid DW, Nicchitta CV. 2015b. LOCAL TRANSLATION. Comment on “Principles of ER cotranslational translocation revealed by proximity-specific ribosome profiling.” Science 348: 1217.

Reimand J, Arak T, Vilo J. 2011. g: Profiler—a web server for functional interpretation of gene lists (2011 update). Nucleic Acids Res gkr378.

Roth FP, Hughes JD, Estep PW, Church GM. 1998. Finding DNA regulatory motifs within unaligned noncoding sequences clustered by whole-genome mRNA quantitation. Nat Biotechnol 16: 939–945.

Rowland AA, Voeltz GK. 2012. Endoplasmic reticulum–mitochondria contacts: function of the junction. Nat Rev Mol Cell Biol 13: 607–625.

Shah P, Ding Y, Niemczyk M, Kudla G, Plotkin JB. 2013. Rate-limiting steps in yeast protein translation. Cell 153: 1589–1601.

Singh G, Kucukural A, Cenik C, Leszyk JD, Shaffer SA, Weng Z, Moore MJ. 2012. The cellular EJC interactome reveals higher-order mRNP structure and an EJC-SR protein nexus. Cell 151: 750–764.

Singh G, Ricci EP, Moore MJ. 2014. RIPiT-Seq: a high-throughput approach for footprinting RNA:protein complexes. Methods 65: 320–332.

Storey JD. 2003. The positive false discovery rate: a Bayesian interpretation and the q-value. Ann Stat 31: 2013–2035.

Sylvestre J, Vialette S, Corral Debrinski M, Jacq C. 2003. Long mRNAs coding for yeast mitochondrial proteins of prokaryotic origin preferentially localize to the vicinity of mitochondria. Genome Biol 4: R44.

Tuller T, Carmi A, Vestsigian K, Navon S, Dorfan Y, Zaborske J, Pan T, Dahan O, Furman I, Pilpel Y. 2010. An evolutionarily conserved mechanism for controlling the efficiency of protein translation. Cell 141 : 344–354.

Valen E, Sandelin A, Winther O, Krogh A. 2009. Discovery of regulatory elements is improved by a discriminatory approach. PLoS Comput Biol 5: e1000562.

Vogel C, Marcotte EM. 2012. Insights into the regulation of protein abundance from proteomic and transcriptomic analyses. Nat Rev Genet 13: 227–232.

Wang S, Yao X. 2012. Multiclass Imbalance Problems: Analysis and Potential Solutions. IEEE Trans Syst Man Cybern B Cybern. http://dx.doi.org/10.1109/TSMCB.2012.2187280.

Wang X, Lu Z, Gomez A, Hon GC, Yue Y, Han D, Fu Y, Parisien M, Dai Q, Jia G, et al. 2014. N6-methyladenosine-dependent regulation of messenger RNA stability. Nature 505: 117–120.

Wang X, Zhao BS, Roundtree IA, Lu Z, Han D, Ma H, Weng X, Chen K, Shi H, He C. 2015. N(6)-methyladenosine Modulates Messenger RNA Translation Efficiency. Cell 161: 1388–1399.

Zinshteyn B, Gilbert WV. 2013. Loss of a conserved tRNA anticodon modification perturbs cellular signaling. PLoS Genet 9: e1003675.

